# Gliding motility of *Plasmodium* merozoites

**DOI:** 10.1101/2020.05.01.072637

**Authors:** Kazuhide Yahata, Melissa N. Hart, Heledd Davies, Masahito Asada, Samuel C. Wassmer, Thomas J. Templeton, Moritz Treeck, Robert W. Moon, Osamu Kaneko

## Abstract

*Plasmodium* malaria parasites are obligate intracellular protozoans that use a unique form of locomotion, termed gliding motility, to move through host tissues and invade cells. The process is substrate-dependent and powered by an actomyosin motor that drives the posterior translocation of extracellular adhesins which in turn propel the parasite forward. Gliding motility is essential for tissue translocation in the sporozoite and ookinete stages; however, the short-lived erythrocyte-invading merozoite stage has never been observed to undergo gliding movement. Here we show *Plasmodium* merozoites possess the ability to undergo gliding motility and that this mechanism is likely an important precursor step for successful parasite invasion. We demonstrate that two human infective species, *P. falciparum* and *P. knowlesi*, have distinct merozoite motility profiles which may reflect distinct invasion strategies. Additionally, we develop and validate a higher throughput assay to evaluate the effects of genetic and pharmacological perturbations on both the molecular motor and complex signaling cascade that regulates motility in merozoites. The discovery of merozoite motility provides a new model to study the glideosome and may facilitate the pursuit of new targets for malaria treatment.

**Significance statement:** *Plasmodium* malaria parasites use a unique substrate-dependent locomotion termed gliding motility to translocate through tissues and invade cells. Dogma has suggested that the small labile invasive stages that invade erythrocytes, merozoites, use this motility solely to penetrate target erythrocytes. Here we reveal that merozoites use gliding motility for translocation across host cells prior to invasion. This forms an important pre-invasion step that is powered by a conserved actomyosin motor and is regulated by a complex signaling pathway. This work fundamentally changes our understanding of the role of gliding motility and invasion in the blood and will have a significant impact on our understanding of blood stage host-pathogen interactions, parasite biology, and could have implications for vaccine development.

## Introduction

Apicomplexan parasites traverse tissues and invade cells via a mechanism known as gliding motility, a process unique to the phylum that uses neither propulsive structures such as flagella or cilia, nor cellular shape changes as for peristaltic and amoeboid motility (1, 2). Motility of invasive forms of malarial parasites (termed “zoites”) was first described for the ookinete stage in avian blood (3), and then for the sporozoite stage in the mosquito (4, 5). Unlike ookinetes and sporozoites, which must traverse through tissues, no gliding motility has been described for the merozoite, which invades erythrocytes in the bloodstream. Instead, only limited reorientation movement and cellular deformation has been observed across several malarial parasite species, including *Plasmodium knowlesi*, *P. falciparum,* and *P. yoelii* (6–8). Due to frequent and passive encounters with erythrocytes in the bloodstream shortly after egress, it has been presumed that merozoites do not require motility, leading to the consensus that the molecular motor is principally required for entry into the erythrocyte (9).

Gliding has perhaps been best described for *Toxoplasma gondii* tachyzoites and *Plasmodium berghei* sporozoites, both of which exhibit helical motion in 3D matrices, as well as during host cell entry (10–12). Treatment of tachyzoites and sporozoites with either cytochalasin D, an inhibitor of actin polymerization, or butanedione monoxime, a myosin inhibitor, stalls both extracellular motility and host cell invasion, indicating that both processes are driven by an actomyosin-based motor (7). Subsequent work discovered that the components of this motor, collectively called the “glideosome”, are located within the space between the parasite inner membrane complex (IMC) and plasma membrane; and are generally conserved between apicomplexan zoites, including merozoites (11, 13, 14). According to the linear motor model, gliding is a substrate-dependent process. Transmembrane ligands are secreted from organelles called micronemes at the apical end of the zoite, where they bind to receptors on the surface of the host cell or tissue via their ectodomains, whilst also connecting to the actomyosin motor through their cytoplasmic tails (11). MyoA engagement with actin subsequently pulls both actin and the receptor-bound ligand rearward, thereby driving the zoite forwards. Finally, the ectodomains of receptor-bound ligands are cleaved by parasite-derived proteases, thus disengaging them from their host cell receptors (11). Deletion of key motor components, including myosin A (MyoA) (15), actin-1 (ACT1) (16), and glideosome-associated protein 45 (GAP45) (17), prevent *P. falciparum* merozoites from invading host erythrocytes, demonstrating the essentiality of the actomyosin motor for erythrocyte invasion (9).

Here we show that both *P. falciparum* and *P. knowlesi* merozoites are capable of gliding motility across both erythrocyte surfaces and polymer coverslips. Distinctive gliding dynamics are found between the two species, suggesting that motility modulates multiple steps of invasion. We developed an assay to evaluate the effect of genetic and pharmacological perturbations on both the molecular motor and complex signaling cascade that regulates motility in merozoites.

## Results

### Gliding motility of *Plasmodium* merozoites

Here we sought to address the long-standing dogma that malarial merozoites do not undergo conventional gliding motility. Whilst motility of sporozoites is normally observed on bovine serum albumin-coated glass slides, merozoites do not glide on this substrate. However, when using polymer coverslips with a hydrophilic coating (ibiTreat), we observed motile merozoites. When imaged immediately after erythrocyte egress, *P. falciparum* merozoites show directional movement on the coverslip surface which displaces them from the hemozoin-containing residual body (Fig. 1*A* and *B*; *SI Appendix,* Movies S*1* and *2*). The average *P. falciparum* merozoite gliding speed was 0.59 μm/second (SD = 0.14 μm/second; n = 10 individual merozoites), considerably slower than that of *P. yoelii* sporozoites (helical gliding 5.02 μm/second; SD = 0.83 μm/second; n = 8), *Toxoplasma gondii* tachyzoites (helical gliding 2.60 μm/second; SD = 0.54 μm/second; n = 13; circular gliding 1.84 μm/second; SD = 0.32 μm/second; n = 13), and *Babesia bovis* merozoites (6.02 μm/second; SD = 0.73 μm/second; n = 5), but significantly faster than *P. berghei* ookinetes (helical gliding 5.8 μm/min) (18). The longest gliding time of *P. falciparum* merozoites was 43 seconds, shorter than those of *P. yoelii* sporozoites (> 600 seconds), *T. gondii* tachyzoites (> 600 seconds) and *B. bovis* merozoites (125 seconds). Thus, the short-lived motility of *P. falciparum* merozoites appears to correlate with their decline in erythrocyte invasion efficiency within a few minutes after egress (19).

**Fig. 1.**
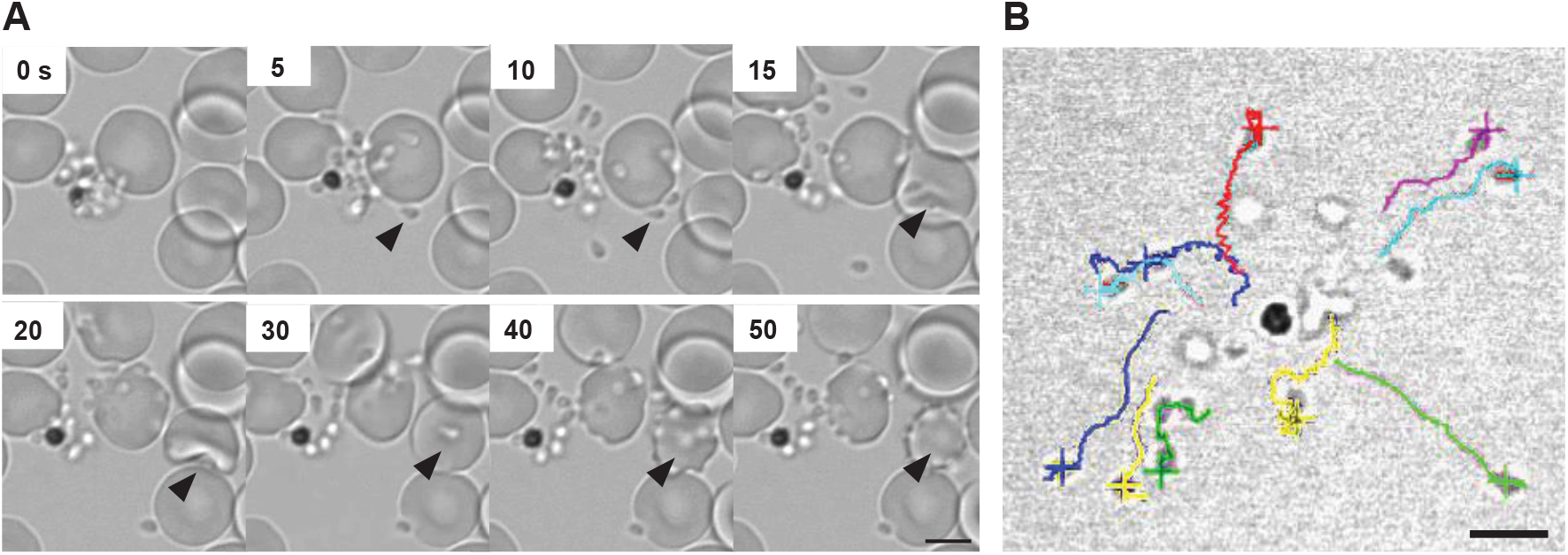
Gliding motility of *P. falciparum* merozoites. (*A*) Time-lapse imaging for *P. falciparum* merozoite gliding motility and erythrocyte invasion. Still images from Movie S1. Arrowhead indicates a merozoite gliding on the coverslip (5 and 10 seconds), followed by erythrocyte deformation (15 and 20 seconds) and merozoite internalisation (30–50 seconds). (*B*) Each merozoite was traced in different colors and gliding speed was evaluated from Movie S2.

The zoonotic malaria parasite, *P. knowlesi,* has much larger and longer-lived merozoites (19–21) and is capable of invading both human and macaque erythrocytes (22); thus, we hypothesized that this may result in different gliding behavior. In the absence of fresh erythrocytes, *P. knowlesi* merozoites also exhibit some motility on ibiTreat coverslips, but the number of motile merozoites increases using poly-L-lysine-coated (PLL) polymer coverslip surfaces (*SI Appendix,* Movie S3). A median 62% (IQR = 55%) of all merozoites within a given schizont exhibited motility on PLL-coated coverslips, with a ‘gliding’ merozoite defined as one which attached to the coverslip surface and exhibited forward movement for at least 5 continuous seconds (Fig. 2*A*). To test whether gliding is surface dependent, *P. knowlesi* merozoites were also monitored on uncoated polymer and glass coverslips. A much lower percentage of motile parasites was observed for the uncoated polymer (mean = 38%; SD = 36%) and glass coverslips (mean = 25%; SD = 22%) (*SI Appendix,* Fig. S1*A*). This suggests that both the coating and the use of polymer rather than glass coverslips is critical for optimal gliding to occur and may account for why merozoite motility has not been observed previously.

**Fig. 2.**
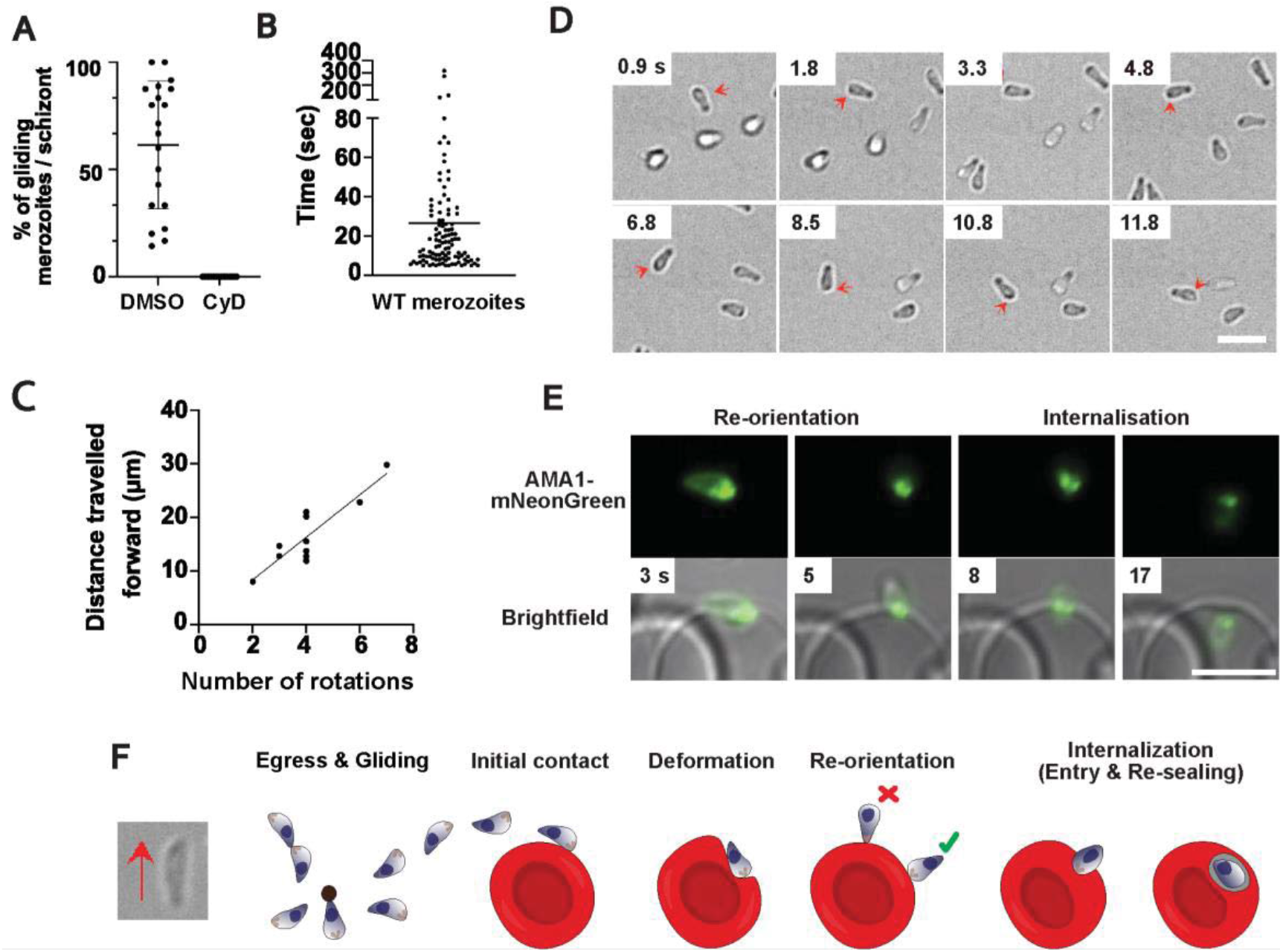
Gliding motility of *P. knowlesi* merozoites. (*A*) The percentage of merozoites within a *P. knowlesi* schizont which exhibit motility, both for DMSO-treated parasites (mean = 62.5%) and 0.1 μM Cytochalasin D (CyD; IC_50_ = 0.023 ± 6.7 nM) -treated parasites (no gliding observed). A ‘motile’ merozoite was defined as having demonstrated directional forward motion along the surface of the coverslip for at least 5 continuous seconds. Each dot is representative of one schizont (n = 20). Error bars denote +/− 1 SD (*B*) The total time each motile *P. knowlesi* merozoite (n = 109; median = 15 seconds) spent gliding during the 10-minute imaging window post-egress. Error bars indicate interquartile range. (*C*) Number of rotations that merozoites completed plotted against the distance travelled for each glide (n = 10). As the number of rotations increased, so did the distance travelled forward, indicating rotation drives forward motion (Pearson correlation coefficient, R = 0.88). (*D*) Time lapse imaging demonstrating a *P. knowlesi* merozoite rotating as it glides. Red arrows indicate a dark spot located to one side of the wider end of the merozoite, which shifts to the opposite side (shown in subsequent frames), as it turns, and then back to the original position to complete a full rotation (see Movie S6). Scale bar represents 5 μm. (*E*) Time lapse imaging depicting a *P. knowlesi* merozoite with mNeonGreen-tagged AMA1 invading an erythrocyte. Panels 1 and 2 demonstrate re-orientation of the wider end of the merozoite to align with the erythrocyte membrane. This is followed by the formation of the moving junction, depicted as two green dots at the merozoite-erythrocyte interface (panel 3), and finally entry into the host cell (panel 4). Scale bar represents 5 μm. (*F*) Schematic illustrating gliding and erythrocyte invasion. Gliding proceeds with the wider, apical end of the merozoite leading. During gliding, merozoites stretch, and a pointed protrusion can be seen at the wider end of the zoite (left hand brightfield image), which engages with the erythrocyte membrane upon re-orientation and internalisation. Re-orientation of the zoite to make a contact of wider end (green tick), and not the thinner end of the zoite as previously hypothesized (red cross), with the erythrocyte membrane occurs prior to entry. During internalisation, constriction of the apical end of the zoite causes the basal end to expand. Finally, after entry is complete, the parasite resides in a parasitophorous vacuole where its development continues.

*P. knowlesi* merozoites were faster (mean = 1.06 μm/second; SD = 0.29 μm/second; n = 57 individual merozoites) than *P. falciparum* (*SI Appendix,* Fig. S1*B*) and were capable of gliding for up to 316 seconds (Fig. 2*B*) on PLL surfaces. To assess the involvement of an actomyosin motor for merozoite motility, egressing merozoites were treated with the actin polymerization inhibitor, cytochalasin D (0.1 μM; IC_50_ = 23 nM). This treatment completely inhibited motility (Fig. 2*A* and *SI Appendix,* Movie S4). Cytochalasin D addition also prevented efficient dispersal post egress and frequently resulted in merozoites remaining attached to each other and/or the residual body (Fig. 2*A* and *SI Appendix,* Movie S4). However, whilst dispersal was inhibited, cytochalasin D treatment did not impede egress itself, as visualized using wheatgerm agglutinin-stained erythrocytes and *P. knowlesi* schizonts treated with the drug (*SI Appendix,* Movie S4). The capacity for erythrocyte egress is in line with observations of cytochalasin D-treated *P. falciparum* parasites (17) (*SI Appendix,* Movie S4). Even without inhibitors, *P. knowlesi* merozoite movement was sometimes impaired by groups of merozoites remaining attached to each other. Notably, though, in many instances, pairs, or smaller groups of merozoites were frequently observed to glide together in unison, when “pointing” in the same direction. Merozoites often completed several glides, with a median cumulative distance of 14 μm (IQR =16 μm), and some travelling as far as 200 μm within the 10-minute imaging window (*SI Appendix,* Fig. S1*D*). The majority of gliding occurred within 5 minutes of egress (*SI Appendix,* Fig. S1*E*), with peak gliding occurring during the initial 1 to 2 minute window. Gliding speed appeared to decline over subsequent glides (*SI Appendix,* Fig. S1*F*), indicative of declining motor function over time, which potentially contributes to the window of merozoite viability. No increase in gliding activity was observed when freshly egressed parasites were first incubated in the dark for 5 minutes prior to commencing imaging for an additional 4 minutes (SI Appendix, Fig. S1*E*). Thus, for *P. knowlesi*, continuous light exposure is unlikely to be the cause of declining motor function (*SI Appendix*, Fig. S1*E*).

Like other apicomplexan zoites (10, 23, 24), *P. knowlesi* merozoites appear to undergo corkscrew-like rotation (*SI Appendix,* Movie S5), with a correlation between the number of turns and forward translocation, indicating a link between the two motions (Fig. 2*C* and *D*). On average, for each body length the merozoite moved forward, it rotated 0.8 times – equivalent to a tangential velocity of 1 μm/second (n = 10 merozoites). This is consistent with a linear motor running at a 42-degree angle down the longitudinal axis of the merozoite. Nine out of ten merozoites rotated counterclockwise, demonstrating the same chirality seen for *Plasmodium* ookinetes (25). Rotation could not be discerned for *P. falciparum* merozoites, likely due to the round morphology and small size.

For both *Plasmodium* species, gliding and invasion proceeded with the wider end of the merozoite leading, rather than the narrower pointed end (Fig. 2*D*). The narrower, pointed end has been widely suggested to contain the apical complex of the parasite, and indeed is consistent with early images of invading parasites by transmission electron microscopy (26). To test whether the apical complex is instead located at the wider end of the parasite, we examined *P. knowlesi* parasites expressing mNeonGreen-tagged apical membrane antigen 1 (AMA1-mNG) using live microscopy. Movie S6 clearly shows that the apical end is located at the wider end of the zoite (Fig. 2*E*), and that host cell entry proceeds in the same orientation as surface gliding, as observed previously for *B. bovis* merozoites (24). Imaging of the AMA1-mNG parasites also shows, for the first time using live microscopy to image *Plasmodium* invasion, the formation of a ring structure of the tight junction as the parasite invades the host erythrocyte (Fig. 2*E* and *SI Appendix,* Movie S6). A small protrusion likely corresponding to the apical complex is visible slightly offset from the apex of the wider front-end (Fig. 2*F*, left hand image). It is the accentuation of this during the constriction of invasion depicted within classic electron microscopy images which has likely led to the general assumption that merozoites uniformly narrow towards the apical end (Fig. 2*F*). Whilst this is most clearly seen in the elongated forms of the *P. knowlesi* merozoites, it is also demonstrated in videos of gliding in *P. falciparum* (*SI Appendix,* Movie S2).

Merozoites also exhibited helical motility during host cell entry and after internalization within the host cell, as has been described for invading *T. gondii* tachyzoites (27). This event can be seen when observing apically localized AMA1-mNG, which rotates around the plane of the moving junction in an anti-clockwise manner during entry and then again rapidly for several seconds post internalization (*SI Appendix,* Movie S6). Thus, like other zoites, *Plasmodium* merozoites appear to enter host cells using a corkscrew-like mechanism. Post entry spinning, as recently visualized with 4D lattice light sheet microscopy for invading *P. falciparum* parasites (28) may, in turn, facilitate twisting and separation of the newly formed parasitophorous vacuole from the erythrocyte plasma membrane.

### Gliding motility is powered by an actomyosin motor and glideosome complex

To determine the characteristics of the *P. falciparum* merozoite glideosome we evaluated the effect of chemical compounds and parasite genetic modifications on merozoite gliding motility. In comparison to *P. knowlesi*, *P. falciparum* merozoites appear to be more vulnerable to light (29) and often lose their ability to invade erythrocytes when exposed to even small amounts of light, complicating the observation of their gliding movement by light microscopy. In our hands, schizonts exposed to light over several minutes often showed very few motile merozoites (*SI Appendix,* Movie S7). On average, only roughly 6% (2/32) of schizonts released 70% or more gliding merozoites, where a ‘glide’ was defined as forward movement for at least 5 continuous seconds (*SI Appendix,* Fig. S3*A* and Movie S2). Thus, to overcome the light sensitivity of *P. falciparum* merozoites we developed an assay in which schizonts were seeded and incubated in the dark on coverslips for 1 hour 37°C until the completion of merozoite egress. Motility could then be quantified by measuring the distance between a DAPI-stained merozoite nucleus and the hemozoin in the residual body (Fig. 3*A*). In comparing with DMSO-treated merozoites (100%), 0.1, 1, and 10 μM cytochalasin (IC_50_ = 0.085 μM) D treatment significantly reduced the merozoite-hemozoin distance (77.9%; IQR = 25.1%, 52.6%; IQR = 28.2%, and 50.9%; IQR = 16.5%, respectively). Treatment with jasplakinolide, an actin filament stabilizer reported to increase the speed of *T. gondii* tachyzoites, did not significantly increase the distance travelled by *P. falciparum* merozoites (Fig. 3*B* and *SI Appendix,* S3).

**Fig. 3.**
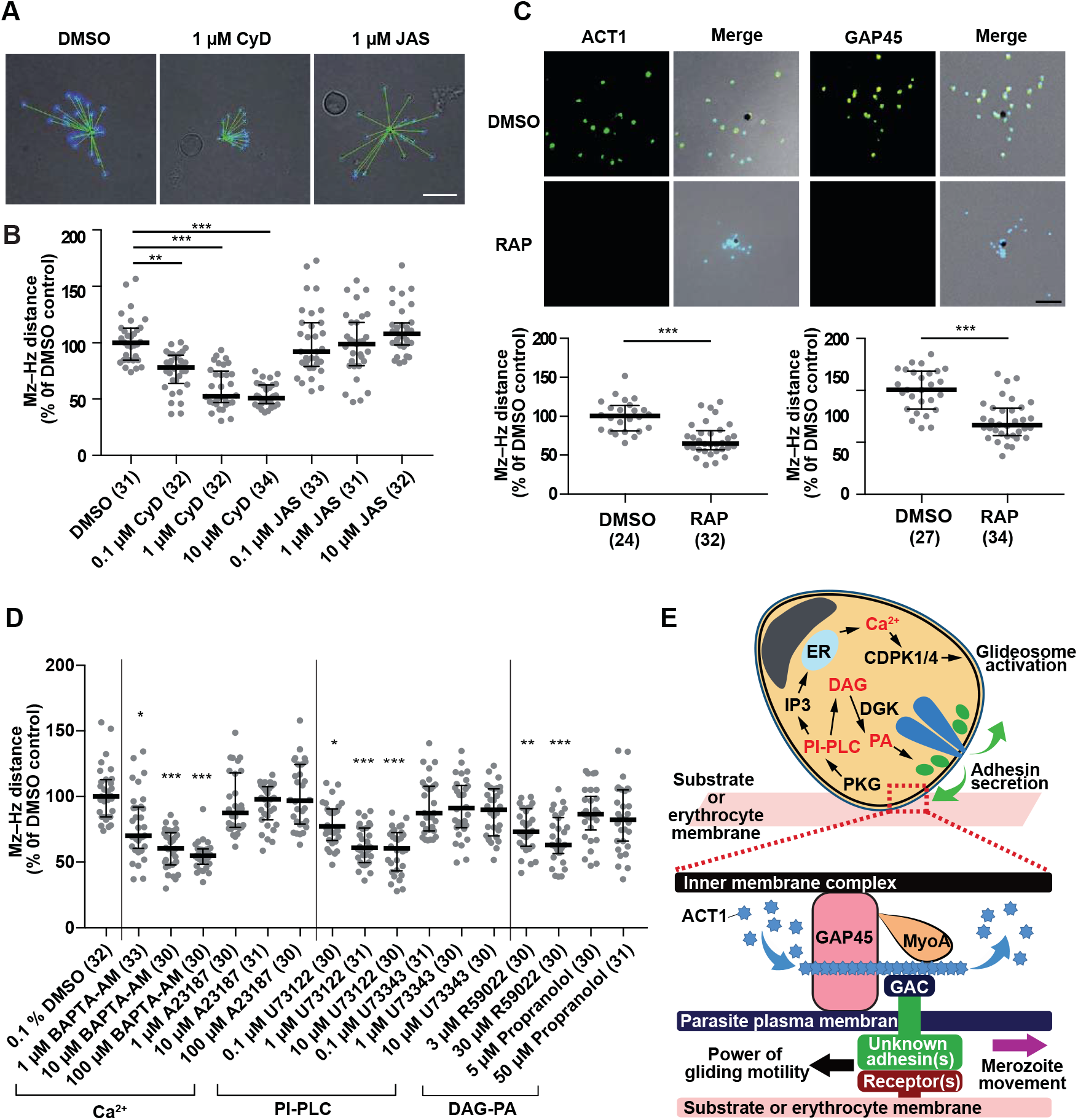
The effects of chemical compounds and parasite genetic modifications on *P. falciparum* merozoite gliding motility. Purified *P. falciparum* schizonts were seeded on the coverslip and merozoites were left to egress for 1 hour. (*A*) The distance of the merozoite nucleus (DAPI, Mz) from hemozoin (black pigment, Hz) was measured (green line, Mz–Hz distance). Where indicated in the y-axes (panels *B* and *C*) the relative Mz–Hz distance compared to DMSO control obtained from each schizont with their median and interquartile range are shown. The number of analyzed schizonts from three independent experiments are indicated in the parentheses. (*B*) Effect of 0.1% DMSO, 0.1, 1, or 10 μM cytochalasin D (CyD, IC_50_ = 0.085 ± 0.029 μM, or jasplakinolide (JAS, IC_50_ = 0.110 ± 0.019 μM) were evaluated for merozoite gliding motility. **, and *** indicate p < 0.001, and < 0.0001, respectively. (*C*) Inhibition of gliding motility in rapamycin (RAP)-treated ACT1- or GAP45-deleted *P. falciparum* parasites. IFA with specific antibodies indicated ACT1 or GAP45 were not detected in RAP-treated transgenic parasites. *** indicates p < 0.0001 by the Mann-Whitney test. (*D*) Purified *P. falciparum* schizonts were treated with BAPTA-AM (IC_50_ = 0.992 ± 0.187 μM), A23187 (IC_50_ = 0.588 ± 0.029 μM), U73122 (IC_50_ = 0.271 ± 0.085 μM), U73343 (IC_50_ = 5.444 ± 0.199 μM), R59022 (IC_50_ = 4.678 ± 0.392 μM), or propranolol (IC_50_ = 0.551 ± 0.135 μM) and merozoite gliding assays were performed. The relative Mz–Hz distance compared to DMSO control obtained from each schizont with their median and interquartile range are shown. The number of analyzed schizonts from three independent experiments are indicated in the parentheses. *, **, and *** indicate p < 0.05, < 0.001, and < 0.0001, respectively. (*E*) Overview of molecular mechanisms for gliding motility of *P. falciparum* merozoite. After merozoite egress from the erythrocyte, merozoite adhesin(s) are secreted from micronemes (green) via a signaling pathway involving phosphoinositide-phospholipase (PI-PLC) and diacylglycerol (DAG) kinase (DGK) and bind to environmental substrates including the erythrocyte membrane. A pathway involving PI-PLC and Ca^2+^ activates calcium dependent protein kinases (CDPKs) and phosphorylates the components of the glideosome machinery (32–35, 58). Grey, nucleus and blue, rhoptries. Gliding motility is powered by an actomyosin motor of the glideosome machinery and the merozoite movement is transferred to the erythrocyte membrane causing erythrocyte deformation upon merozoite attachment. ACT1, actin-1; IMC, inner membrane complex; PKG, cyclic GMP-dependent protein kinase; PA, phosphatidic acid; GAP45, glideosome-associated protein 45; MyoA, myosin-A; and GAC, glideosome-associated connector.

We next examined the effects of the conditional deletion of two essential glideosome components, ACT1 (16) and GAP45 (17). Transgenic lines were able to egress after both the control DMSO treatment and upon rapamycin induced gene excision, but the merozoite–hemozoin distance was significantly reduced after deletion of either ACT1 or GAP45 (Fig. 3*C*). When AMA1, a microneme protein important for erythrocyte invasion but unlikely to be involved in merozoite motility (30, 31) was conditionally deleted, parasites were able to efficiently egress and move across the substrate (*SI Appendix*, Fig. S4). These results confirm the requirement of the glideosome for *Plasmodium* merozoite gliding motility.

### Gliding motility is regulated by a complex signaling pathway

Microneme discharge plays an essential role in the egress, gliding motility, and cell invasion of apicomplexan parasites and is regulated by a set of intracellular signaling enzymes, including calcium-dependent protein kinases (32, 33), phosphoinositide-phospholipase C (PI-PLC) (34), and diacylglycerol (DAG) kinase (35). We evaluated whether these enzymes are also involved in the gliding motility of *P. falciparum* merozoites. Although the calcium ionophore A23187 (up to 100 μM) did not show a significant effect, the calcium chelator BAPTA-AM (10 μM) significantly reduced merozoite–hemozoin distance (p < 0.0001; Fig. 3*D* and *SI Appendix,* S3*B*). The PLC inhibitor U73122 (1 μM), but not the inactive analog U73343 (up to 10 μM), significantly reduced merozoite–hemozoin distance (p < 0.0001). The DAG kinase inhibitor R59022 (3 μM), which inhibits the conversion of DAG to phosphatidic acid (PA) also significantly reduced movement (p < 0.001); while the merozoite–hemozoin distance was not changed with propranolol, an inhibitor of phosphatidate phosphohydrolase (the converter of PA to DAG). Collectively, these results are consistent with reports on *Toxoplasma* tachyzoites (35) and indicate that complex signaling pathways are involved in gliding motility of *P. falciparum* merozoites (Fig. 3*E*).

### Gliding motility facilitates erythrocyte-specific interactions and underpins important precursor steps to invasion

Having established the characteristics of merozoite motility and its regulation, we next sought to understand whether gliding may play a role during erythrocyte invasion in addition to host cell entry, and whether this function might also facilitate movement of merozoites across alternative cellular surfaces. Prior to host cell entry, merozoite-erythrocyte interactions cause the erythrocyte membrane to deform, often resulting in the entire erythrocyte “wrapping” around the merozoite (7). Several studies have highlighted the importance of deformation as a mechanism merozoites use to commit to host cell entry and have demonstrated that the merozoite’s molecular motor is required for deformation (16, 17, 36). However, the precise mechanism behind this phenomenon has not been elucidated. In good agreement with previous work, we found that not only were rapamycin-treated ACT1- or GAP45-deleted *P. falciparum* parasites unable to glide, but they were also unable to deform erythrocytes (Fig. 4*A* and *B*). Thus, we reasoned that gliding motility itself may be required for erythrocyte deformation.

**Fig. 4.**
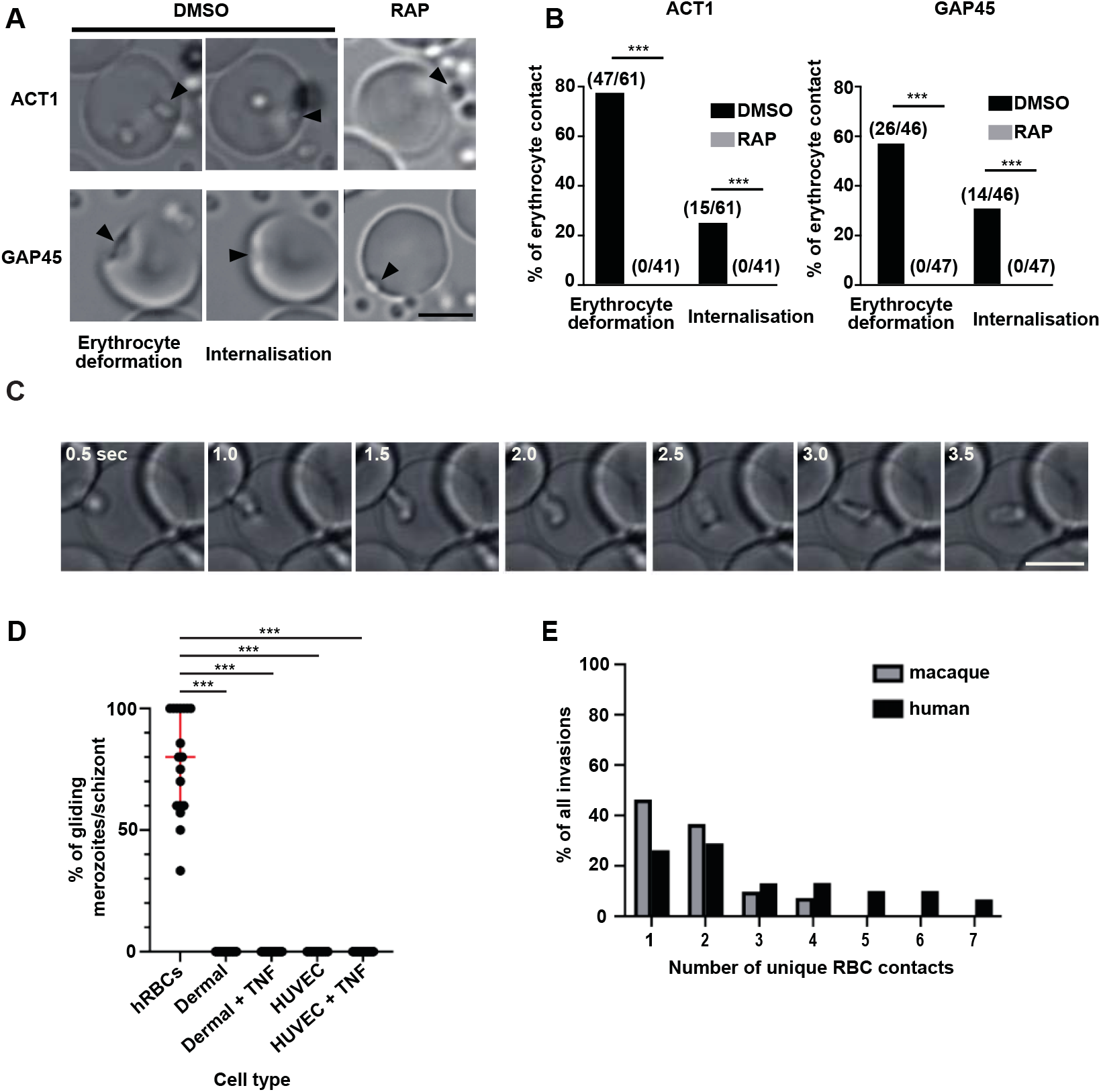
A role for gliding motility in facilitating merozoite-erythrocyte interactions. (*A* and *B*) Erythrocyte deformation and merozoite internalisation events were seen for DMSO-treated ACT1- or GAP45-floxed *P. falciparum* parasites, but not detected after RAP-treatment (p < 0.001 for all by two-tailed Fisher’s exact test). Scale bar represents 5 μm. (*C*) Still images taken from Movie S7, depicting a *P. knowlesi* merozoite beginning to deform a human erythrocyte as gliding motility is initiated along the surface of the host cell. Scale bar represents 5 μm. (*D*) A median 80% of merozoites within a given schizont demonstrate gliding motility along human erythrocytes (n = 18 schizonts). In contrast, no gliding is observed at all on either TNF stimulated or non-stimulated HUVEC (n = 16 and 10 schizonts, respectively) and dermal endothelial cells (n = 16 and 10 schizonts, respectively). Comparisons were made by one-way ANOVA, using a Kruskal-Wallis test; *** indicates p<0.0001. (*E*) The total number of unique erythrocyte contacts each invading merozoite made was significantly lower for macaque (n = 41 invasions) vs human (n = 38 invasions) erythrocyte invasions. Comparisons were made using Poisson regression (Poisson coef. 0.076, 95% CI - 0.37 - 0.52; P<0.001).

Due to our finding that the majority of *P. knowlesi* merozoites do not disperse from each other when gliding is inhibited with cytochalasin D, we could not easily measure the impact of gliding inhibition on erythrocyte deformation for this species. However, the larger size and more pronounced morphology of *P. knowlesi* merozoites did allow us to visualize the deformation process of wild type merozoites more clearly. Deformation did not appear to simply be the result of the merozoite attempting to be internalized in the erythrocyte, as previously hypothesized (6). Rather, deformation was marked by a wrapping of the erythrocyte membrane down the length of a forward moving merozoite, horizontally aligned with the erythrocyte surface. Notably, deformation began as a merozoite initiated gliding across a host cell surface, with waves of deformation often travelling from the zoite’s apical prominence to the basal end (Fig. 4*C* and *SI Appendix,* Movie S8). These sum results indicate that not only does gliding motility coincide with erythrocyte deformation, but also that the two processes are mechanically linked.

Recently, both sexual and asexual stages of multiple *Plasmodium* species have been identified within bone marrow compartments (37–39). This raised the question of whether merozoites may be able to glide on endothelial surfaces and/or within extravascular compartments, such as the bone marrow. Yet how do *Plasmodium* parasites exit the bloodstream and enter tissues? One possibility is that parasitized reticulocytes may be returning to extravascular environments via extravasation. However, another mechanism could be that free merozoites are capable of gliding across endothelial barriers and into tissues (40, 41). To test the latter theory, we examined *P. knowlesi* parasites egressing on either primary human umbilical vein (HUVEC) or dermal endothelial cell lines. Rather surprisingly, we found that *P. knowlesi* merozoites were unable to adhere to or demonstrate motility across either cell line (Fig. 4*D* and *SI Appendix,* Movie S9). Similar results were obtained when endothelial cells were pre-treated with tumor necrosis factor (TNF), an inflammatory cytokine known to increase the expression of endothelial adhesive surface ligands (Fig. 4*D*) (42). Thus, whilst these results do not rule out the possibility of gliding within tissue compartments, they do suggest that gliding is unlikely to support extravasation of free merozoites and that merozoite motility may be specific to erythrocyte lineages and highly adherent artificial surfaces, in strong contrast to other zoites which can glide successfully on a broad array of substrates.

Despite the finding that merozoites are unlikely to use gliding to move across endothelial barriers, we did observe that freshly egressed *P. knowlesi* merozoites often glide across several different human erythrocytes prior to invading their host cell of choice (*SI Appendix,* Movie S10). Thus, we reasoned that in addition to facilitating erythrocyte deformation, gliding may enable merozoites to contact multiple erythrocytes, in the event they are unable to invade the first host cell they encounter. Studies have shown that most invading *P. falciparum* merozoites (∼85%) will invade the first cell they contact (36); however, we found that only 26% (10/38 invasions) of invading *P. knowlesi* merozoites invaded the first human erythrocyte they contacted, and a significant proportion of merozoites (∼32%, or 12/38 events) contacted 4 to 7 unique cells prior to committing to invasion (Fig. 4*E*). We hypothesized that this stark difference between species may be because, unlike *P. falciparum*, the natural host cells of *P. knowlesi* are not human, but rather macaque erythrocytes (22). Thus *P. knowlesi* merozoites may need to contact more human erythrocytes overall, to encounter an erythrocyte amenable to invasion. In support of this theory, we found that on average, *P. knowlesi* merozoites contacted significantly fewer macaque erythrocytes prior to invasion (Fig. 4*E*), with the vast majority of merozoites (85%) invading either the first or second macaque cell they contacted. Therefore, these results indicate that gliding could contribute to the zoonotic potential of *P. knowlesi* and may explain how merozoites of multiple species navigate strict erythrocyte tropism requirements to proliferate under sub-optimal conditions.

## Discussion

In this work, we show for the first time that *Plasmodium* merozoites undergo gliding motility and demonstrate productive movement across coverslip and erythrocyte surfaces for two human infective species: *P. falciparum* and *P. knowlesi*. Motility was more evident for *P. knowlesi* merozoites, which in our hands were found to glide on average nearly twice as fast and for 7 times longer than *P. falciparum* merozoites.

Since the first, and landmark video microscopy of *P. knowlesi* in 1975 (6), no further imaging studies have been published for this species. This, in combination with the frequent use of glass slides to image parasites, rather than the polymer coverslips that were used for this study, may explain in part why motility was not observed in *P. falciparum* invasion studies. However, our retrospective analysis of published data shows several subtle examples of *P. falciparum* merozoites completing short glides across human erythrocytes, such as videos from Treek et al., 2009 (Video S4, from 25-70 seconds) (30) and Weiss et al., 2015 (Video S13, from 0-26 seconds) (36).

Our work, in good agreement with previous studies (16, 17, 36), shows that without motor function merozoites cannot glide across cellular surfaces or deform host erythrocytes. Thus, gliding motility is likely an important requirement for deformation. While deformation itself is not an essential invasion step for *P. falciparum*, strong deformation has been positively correlated with invasion success and is predicted to increase the efficiency of the host cell selection process (36). Therefore, by extension, gliding is also likely to facilitate these key steps, increasing the likelihood of each merozoite-erythrocyte contact progressing into invasion.

Having clearly demonstrated gliding motility in static in vitro cultures, it will be important to determine how this mechanism translates to flow conditions found *in vivo*. Whilst the feasibility of cell-cell transfer is yet to be tested under flow conditions, live *in vivo* imaging of erythrocytes flowing through human capillaries shows that erythrocytes are frequently attached to each other for prolonged periods of time, travelling together at roughly the same speed (43). Therefore, in theory, flowing erythrocyte “bridges” may facilitate merozoite movement from cell to cell *in vivo*.

Erythrocytes are known to be highly heterogeneous, in terms of maturity, surface receptor density, and membrane tension properties (44). As such, motility could also enable merozoites to contact a wider range of erythrocytes in the bloodstream, whereby the parasite moves around the surface of an individual erythrocyte, or across the surfaces of multiple erythrocytes until it is able to engage with a critical threshold of invasion receptors or select a cell with favorable characteristics for invasion. Our work has shown that only 26% of *P. knowlesi* merozoites invade the first human erythrocyte they contact, with the vast majority instead contacting multiple cells before committing to invasion. Notably, *P. knowlesi* exhibits a preference (not restriction) for younger erythrocytes when cultured with human erythrocytes (45, 46). Thus, gliding may be a critical mechanism *P. knowlesi* merozoites use to successfully proliferate in human hosts.

A greater proportion of *P. knowlesi* merozoites will invade the first macaque erythrocyte they contact (∼ 46%), in line with their preference for their natural host cells (macaque erythrocytes) over human cells (45). In contrast, most *P. falciparum* merozoites (∼86% of 3D7 parasites) invade the first human erythrocyte they encounter (36), therefore, the ability to contact multiple cells is likely to be less important for *P. falciparum*, and instead gliding may simply enhance erythrocyte receptor interactions on any given cell.

One reason for these distinct strategies could be that each species possesses highly divergent sets of RBL/Rh and DBP/EBA ligands - two critical families of invasion ligands which are key determinants of host cell tropism and are predicted to facilitate host cell selection (36, 47). Differing expression profiles, along with differing levels of receptor abundance and availability within erythrocyte populations, likely impact how efficiently merozoites can “commit” to invasion, and thus their need to contact different cells. At one extreme, for *P. vivax* a single DBP ligand and selection of RBL ligands restricts this species to growth in Duffy positive human reticulocytes (48, 49). Since reticulocytes make up less than 2% of red blood cells circulating within the bloodstream (50), gliding motility may enable *P. vivax* merozoites to roam and eventually encounter reticulocytes.

It is also plausible that motility supports translocation and invasion within tissues such as the bone marrow, which is known to be a significant parasite reservoir for *P. vivax* (39). Parasites may accumulate within the bone marrow via extravasation: the crossing of either infected reticulocytes or free merozoites into the bone marrow parenchyma via a layer of endothelial cells lining the sinusoidal capillaries (40, 41). However, we did not observe merozoite motility across endothelial surfaces; an outcome which suggests that gliding may not support extravasation of free merozoites. Notably, though, *P. vivax* merozoites have been shown to exhibit a preference for the CD71-positive youngest reticulocytes found predominantly within the bone marrow prior to further maturation and intravasation (40). This suggests that at least some invasions may take place within the bone marrow parenchyma itself, contributing to the overall parasite burden within this tissue. Thus, gliding may enable merozoites that have egressed within the bone marrow to specifically target these young reticulocytes in this richly diverse cellular environment.

Our work also enabled us to reverse the dogma perception of the morphology of merozoites, with clear evidence from both gliding and fluorescently tagged parasites demonstrating that the apical complex resides in a small protrusion in the wider end of the zoite, rather than the pointed end of a teardrop shape as it is often depicted (51). *Plasmodium* ookinetes also lead with their wider end (18), and thus it must be reevaluated how we interpret images of invasion and understand the biophysical processes involved (51).

Our displacement assay also enabled us to test the impact of various inhibitors of calcium signaling pathways, previously shown to be important for microneme secretion (34) and invasion (34, 36) for *Plasmodium* parasites. Our results are in good agreement with data from these studies; however, ascribing function based on inhibitor data alone can be challenging. For instance, although our work shows that the inhibitor U73122, but not its inactive analog, U73343, inhibits signaling pathways also predicted to be required for motility; in other systems, the effects of both compounds have been shown to be identical, suggesting off-target activity (52). Therefore, future work should include targeted genetic approaches such as conditional knockout studies ablating signaling enzymes – something which is now readily achievable for *Plasmodium* blood stages in both *P. knowlesi* and *P. falciparum* (53).

One of the outstanding questions is which parasite proteins are required for bridging the motor to host cell ligands to enable gliding motility. Apicomplexan zoites utilize type I transmembrane proteins belonging to the TRAP family to adhere to environmental substrates for gliding. Two such proteins, merozoite thrombospondin-related anonymous protein (MTRAP) and thrombospondin-related apical membrane protein (TRAMP or PTRAMP), are expressed at the merozoite stage (54). MTRAP is dispensable for *P. falciparum* and *P. berghei* merozoites (55); however, multiple studies indicate that TRAMP is essential for the blood stage parasite (56, 57), making it a prime candidate for future work to identify a merozoite gliding adhesin. Our gliding assays can greatly facilitate the identification of such proteins and their roles during invasion.

In conclusion, *Plasmodium* merozoites have the capacity for gliding motility, powered by a conserved actomyosin motor and glideosome complex, and controlled by a complex signaling cascade. The distinct gliding profiles of two different human infective species suggest divergent invasion strategies which provide new mechanisms to address questions of host selectivity and tissue reservoirs of the erythrocytic stages.

## Materials and Methods

### Parasite culture and transfection

*P. falciparum* Dd2 parasites were maintained with O^+^ human erythrocytes in RPMI1640 medium (Invitrogen, CA, USA) supplemented with 25 mM HEPES (Sigma, St. Louis, USA), 0.225% sodium bicarbonate (Invitrogen), 0.1 mM hypoxanthine (Sigma), 25 μg/mL gentamicin (Invitrogen), 0.5% AlbuMax I (Invitrogen), essentially as described (59). The ACT1 (16), GAP45:loxP (17), and AMA1:loxP *P. falciparum* lines (60) were cultured with A^+^ human erythrocytes. WR99210 and G418 were used to generate ACT1 and AMA1:loxP parasite lines, respectively. *P. yoelii* 17XNL parasites were intravenously inoculated into ICR mice (SLC Inc., Shizuoka, Japan). Animal experiments were approved by the Animal Care and Use Committee of Nagasaki University (Permit number: 1403031120-5). The *T. gondii* RH strain was cultured in a confluent monolayer of human foreskin fibroblasts (HFFs) maintained in Dulbecco’s Modified Eagle Medium (DMEM), GlutaMAX supplemented with 10% fetal bovine serum, at 37°C and 5% CO_2_. The *B. bovis* Texas strain was maintained in purified bovine erythrocytes with GIT medium (WAKO, Osaka, Japan) at 37°C with a microaerophilic stationary-phase culture system. A1-H.1 *P. knowlesi* parasites were maintained in human erythrocytes (UK National Blood Transfusion Service) with custom made RPMI-1640 medium, supplemented with 10% Horse Serum (v/v) and 2 mM L-glutamine according to previously described methods (22). Mature schizonts were purified by gradient centrifugation on a 55% Nycodenz layer (Progen, Heidelberg, Germany), as described (22). Tightly synchronized schizonts were transfected using the Amaxa 4-D electroporator and P3 Primary Cell 4D Nucleofector X Kit L (Lonza) according to the protocol described by Moon *et al*. (22).

### Generation of *P. knowlesi* parasites with mNeonGreen-tagged AMA1

*P. knowlesi* parasites with mNeonGreen-tagged AMA1 were generated by insertion of an mNeonGreen sequence immediately before the *AMA1* stop codon (*SI Appendix,* Fig. S2*A*) using the CRISPR-Cas9 system described by Mohring *et al.* (61) (sgRNA sequence: GAGAAGCCTTACTACTGAGT). Donor DNA was synthesized by overlapping PCR, as previously described for PkAMA1-HA tagged parasites (61) and included the mNeonGreen sequence flanked by 500 bp sequences homologous to the C-terminal (HR1) and 3’UTR (HR2) regions of the AMA1 locus (*SI Appendix,* Fig. S2*A*). Primers for PCR are listed in Table S1. In brief, HR1 and HR2 were both PCR-amplified from *P. knowlesi* A1-H.1 gDNA (with primers P6/P7 and P8/P9, respectively), while the mNeonGreen sequence was amplified from plasmid Pk_mNeonGreen with primers P10/P11. All three fragments were subsequently assembled together in two successive steps: firstly by fusing fragments HR1 and mNeonGreen (primers P12/P13), and secondly by fusing fragments HR1/mNeonGreen and HR2 (primers P12/P15) to create the final product, HR1/mNeonGreen/HR2. Post transfection, integration of donor DNA was confirmed by diagnostic PCR, using primers P1 and P3 (*SI Appendix,* Fig. S2*B*). Expression of the AMA1-mNeonGreen fusion protein was also confirmed by indirect immunofluorescence assay (*SI Appendix,* Fig. S2*C*). Air-dried smears of late stage schizonts were fixed in 4% paraformaldehyde for half an hour and permeabilised with PBS containing 0.1% Triton-X100 for 10 minutes. Slides were subsequently blocked overnight in PBS containing 3% BSA before labelling with anti-mNeonGreen mouse monoclonal antibody (32F6; 1:300; Chromotek, Germany) followed by Alexa Fluor 488 goat anti-mouse antibody (1:1000, Invitrogen). Nuclei were stained with DAPI (Invitrogen) and mounted with ProLong Gold antifade reagent (Invitrogen). Images were collected using an inverted microscope (Ti-E; Nikon, Japan) with a 60x oil objective lens (N.A. 1.4).

### Inducible gene-knockout *P. falciparum* parasites

The *GAP45*, *Act1*, and *AMA1* genes were excised by rapamycin treatment from GAP45:loxP, ACT1, and AMA1:loxP *P. falciparum* parasites, respectively (62). Briefly, ring stage parasites synchronized by 5% sorbitol method were treated with 100 nM rapamycin (Sigma) or 0.1% DMSO for 12 hours. Schizonts were purified with a CS magnet separation column (MACS, Miltenyi Biotech, Germany) and used for gliding or erythrocyte invasion assays.

### Endothelial cell culture and stimulation

Commercially available primary human dermal microvascular endothelial cells (HDMVECs, ScienCell, Carlsbad, CA) and umbilical vein endothelial cells (HUVECs, ScienCell) were cultured at 37°C with 5% CO_2_ on fibronectin-coated 6-channel μ-Slides (ibidi, Germany) and grown to confluence in DME/F12 medium (Thermo Fisher, Waltham, MA), pH 7.4, supplemented with 10% fetal bovine serum (ScienCell), 30 μg/mL endothelial cell growth supplement (Sigma), and 10 μg/mL gentamycin (Thermo Fisher). Cells were rinsed with sterile PBS (Sigma) and fresh medium was added 18 hours prior to the gliding assays, with or without 10 pg/mL recombinant human tumor necrosis factor alpha (TNFα, PeproTech, Rocky Hill, NJ). Primary HDMVECs and HUVECs were used between passage 2 and 3 for all experiments.

### Time lapse imaging for the gliding motility

Time lapse imaging assays for *P. falciparum* merozoites were performed at 37°C using an inverted microscope (Ti-E; Nikon) with a 60x oil objective lens (N.A. 1.4 or 1.47). *P. falciparum* synchronized-schizonts in incomplete medium without AlbuMAX I were transferred to the ibiTreat μ-Slide I^0.4^ Luer channel slide (ibidi) and incubated for 10 minutes at 37°C to allow the parasite-infected erythrocytes to attach to the bottom. Incomplete medium was removed and replaced with complete RPMI medium prewarmed to 37°C, then parasites were observed by microscopy. Likewise, synchronized *P. knowlesi* schizonts were transferred using the same technique to either ibiTreat, poly-L-lysine-coated, uncoated, or glass μ-Slide I^0.4/0.5^ Luer channel slides (ibidi) in incomplete RPMI medium and incubated at 37°C for 10 minutes to allow cell attachment. Subsequently, incomplete medium was replaced with complete RPMI medium with 10% horse serum, as per normal culturing conditions. For the actin inhibitor treatments, *P. knowlesi* schizonts were allowed to attach to coverslips while suspended in incomplete RPMI medium, which was then replaced with complete RPMI medium additionally containing 0.1 μM cytochalasin D (Sigma) or 0.1% DMSO (Sigma). For experiments visualizing *P. knowlesi* egress, schizonts were first incubated with 10 μg/mL wheatgerm agglutinin for 10 minutes at 37°C. Schizonts were subsequently washed 2 times in RPMI medium before loading into channel slides for drug treatment as described above. For *P. knowlesi* invasion assays, purified schizonts were added to fresh human (UK National Blood Transfusion Service) or cynomolgus macaque (Silabe) erythrocytes to make a 5-10% parasitaemia and 2.5% haematocrit culture. The haematocrit was subsequently adjusted to 0.25%, and 150 μL culture was loaded into a PLL-coated μ-Slide VI 0.4 (ibidi). *P. yoelii* sporozoites were isolated in RPMI medium from parasite infected mosquito midguts as described (63), then isolated sporozoites were transferred to an ibiTreat μ-Slide I^0.4^ Luer channel slide. *T. gondii* tachyzoites growing in HFFs were collected by scraping after the culture medium was replaced with ENDO buffer (64). Intracellular parasites were isolated from HFFs by lysing host cells via passaging 20 times through a syringe and tachyzoites were transferred to an ibiTreat μ-Slide I^0.4^ Luer channel slide and incubated for 15 minutes at 37°C. The slide was placed on the microscope stage, and the medium was replaced with DMEM before observation. *B. bovis* parasites were isolated in RPMI medium then transferred to the ibiTreat μ-Slide I^0.4^ Luer channel slide. All parasites were observed by differential interference contrast or bright field at 1.5V/100W of halogen lamp or LED light (pT-100; CoolLED, UK) to minimize cell damage. Time-lapse images were captured at 1–100 frames per second using a digital camera (ORCA-R2 or ORCA-Flash4.0; Hamamatsu photonics, Shizuoka, Japan) and imaged using the NIS-Element Advanced Research imaging software (Nikon). The speed of individual merozoites was calculated by tracking actively moving merozoites either manually, using distance measurement tools, or by the tracking module within the NIS-Element software (Nikon). The tangential speed of *P. knowlesi* merozoites was determined by calculating the number of rotations/minute and multiplying this value by the average circumference of a merozoite. The angle of the motor was subsequently calculated using the formula Tan(x) = R/L, where x = the angle of the motor, R = the average distance each merozoite rotated/per body length travelled forward, and L = the body length of the merozoite.

### *P. falciparum* merozoite gliding assay

*P. falciparum* schizonts were purified with a CS magnet separation column, then adjusted to 1 x 10^5^ cell/mL with incomplete RPMI medium and loaded onto an ibiTreat μ-Slide VI^0.4^ channel slide (ibidi). The channel slides were incubated for 10 minutes at 17°C to allow schizont attachment to the bottom followed by replacing the medium with complete RPMI medium containing chemical compounds or DMSO control. Slides were incubated at 17°C for 1 hour then the temperature was increased to 37°C for 1 hour to allow parasite egress. Parasites were fixed with 1% paraformaldehyde fixation solution, which was then replaced with PBS containing 3% BSA (Sigma) and 100 ng/mL DAPI. For the indirect immunofluorescence assay, parasites were fixed in 4% paraformaldehyde containing 0.0075% glutaraldehyde (Nacalai Tesque, Japan) and permeabilized with PBS containing 0.1% Triton-X100 (Calbiochem, CA, USA), then blocked with PBS containing 3% BSA. Next, samples were immunostained with mouse anti-*P. falciparum* ACT1 (final dilution 1:500; a kind gift from Jake Baum) or rat anti-HA (1:1000; Roche, Basel, Switzerland) for HA-tagged GAP45 and AMA1. This was followed by 3×washes with PBS then incubation with Alexa Fluor 488 goat anti-mouse or Alexa Fluor 594 goat anti-rat antibodies (1:1000; Invitrogen) in PBS containing 3% BSA with DAPI. Stained parasites were mounted with Prolong Gold antifade reagent. Microscopy images (Ti-E, Nikon) of egressed merozoites were cropped to 47 x 47 μm^2^ to measure the distance of merozoite nuclei (stained with DAPI) from hemozoin in the residual body (malaria pigment, with bright field image) using NIS-Elements software (Nikon). Statistical analysis was performed by the Kruskal-Wallis test followed by Dunn’s multiple comparison test using PRISM 6 software (GraphPad Software, Inc., CA, USA).

### Chemical Compounds

Complete RPMI medium was supplemented with cytochalasin D, jasplakinolide (Sigma), 1,2-Bis(2-aminophenoxy)ethane-N,N,N’,N’-tetraacetic acid tetraacetoxymethyl ester (BAPTA-AM, Invitrogen), calcium Ionophore A23187 (Sigma), U73122 (Calbiochem), U73343 (Calbiochem), R59022 (Tocris bioscience, UK), propranolol (Sigma), or DMSO. Compound concentrations were as described (33, 34). IC_50_ values for *P. falciparum* were determined using a protocol available at WorldWide Antimalarial Resistance Network (WWARN-http://www.wwarn.org/sites/default/files/INV08_PFalciparumDrugSensitivity.pdf).

## Acknowledgements

The authors thank Sujaan Das and Markus Meissner (supplying *P. falciparum* ACT1), Abigail Perrin and Michael Blackman (supplying *P. falciparum* GAP45:loxP), Alex Hunt (maintaining *T. gondii*), Nattawat Chaiyawong and Edwin Too (maintaining *P. yoelii*). We also thank Reiko Tanaka, Nana Matsumoto, and Momoko Sakura for technical assistance. We are grateful to Japanese Red Cross Blood Society and UK NHS Blood and Transfusion Service for providing human erythrocyte and plasma. This study was conducted at the Joint Usage/Research Center on Tropical Disease, Institute of Tropical Medicine, Nagasaki University, Japan and London School of Hygiene and Tropical Medicine, UK. This work was supported by Fund for the Promotion of Joint International Research, Fostering Joint International Research, 16KK0183 (KY), 19KK0201 (KY), MEXT, Japan. This work was also supported in part by the Grants-in-Aids for Scientific Research, 15K08448 (KY), 19K07525 (KY), 16H05184 (OK), and 19H03461 (OK), MEXT, Japan. RWM was supported by a UK Medical Research Council Career Development Award (MR/M021157/1) and MNH was supported by a Bloomsbury Colleges Studentship. HD and MT receive funding from The Francis Crick Institute, which receives its core funding from Cancer Research UK (FC001189), the UK Medical Research Council (FC001189), and the Wellcome Trust (FC001189). The funders had no role in study design, data collection and analysis, decision to publish, or preparation of the manuscript.

## Author Contributions

KY, MNH, RWM, SCW, MA, MT, and OK conceived and designed research. KY and MNH performed research. HD contributed the generation of transgenic parasite lines. SCW and MNH designed and performed the endothelial surface gliding assays. KY, MNH, TT, MT, RWM, and OK wrote the paper, and all authors contributed to the manuscript and analyzed the data.

## Data and materials availability

All data are available in the manuscript.

## Supplemental Information

This PDF file includes:

Figures S1 to S4

Tables S1

Legends for Movies S1 to S10

Other supplementary materials for this manuscript include the following:

Movies S1 to S10

### Supplementary Figures

**Fig. S1.**
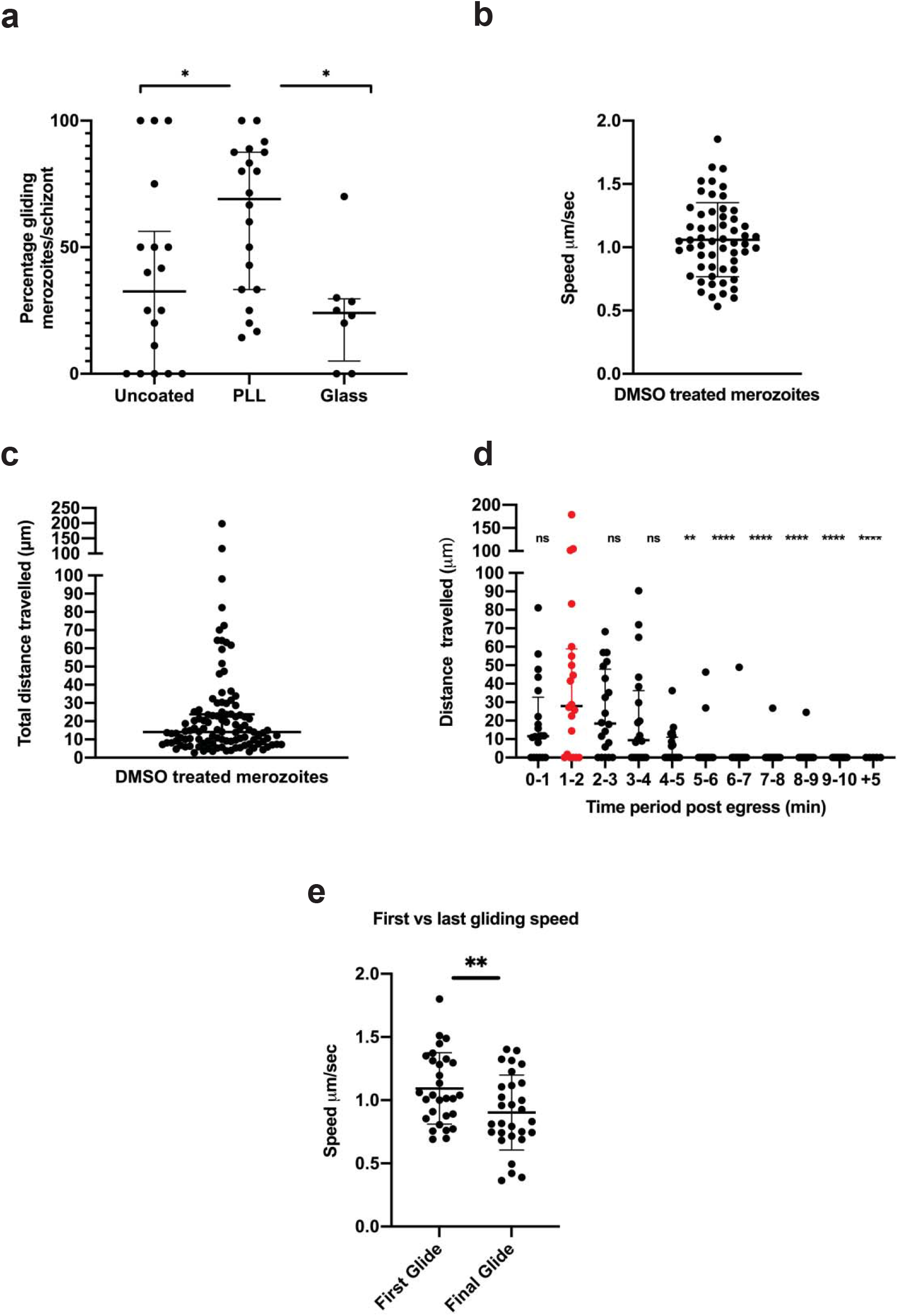
*Plasmodium knowlesi* merozoite gliding speed and duration. **a,** merozoite motility on different surfaces. The percentage of merozoites exhibiting motility decreases from 62% on poly-L-lysine surfaces (n = 20 schizonts) to 38% on uncoated surfaces (n = 18 schizonts; * p < 0.05) and 25% on glass surfaces (n = 8; * p < 0.02). Means compared using one-way ANOVA and Dunnett’s multiple comparison test. Error bars denote +/− 1 s.d. **b,** Speeds of individual merozoites (average = 1.06 μm/second; n = 57 merozoites). Error bars denote +/− 1 s.d. **c**, Total distance travelled by each merozoite. Merozoites travelled a median distance of 14 μm during the 10-minute window of imaging (minimum = 2.8 μm, maximum = 198.6 μm; n = 109 merozoites). **d**, Distances travelled by schizonts (a total of distances travelled by each merozoite) during each minute post egress (n = 20 schizonts) as well as after a 5-minute incubation in the dark (post egress) prior to commencing filming for 3 further minutes (designated as “+5”; n = 5 schizonts). The majority of gliding occurred within 5 minutes post egress, with peak gliding (median of 28 μm travelled) occurring 1-2 minutes post egress (** p < 0.01, **** p < 0.0001, and ns, not significant, as determined by a Kruskal-Wallis test). This delay is likely due to a small ‘settling period’ during the first 60 seconds, while merozoites disperse and begin to connect to the slide coverslip. Error bars denote interquartile range. **e**, First vs last gliding speeds. Comparison by two-tailed paired t-test between the speed of the first and final glides of merozoites (** p < 0.005; n = 29) shows that gliding speed decreases from 1.09 μm/second (average first glide) to 0.90 μm/second (average last glide), indicative of decreasing gliding efficiency over time. Error bars denote +/− 1 s.d.

**Fig. S2.**
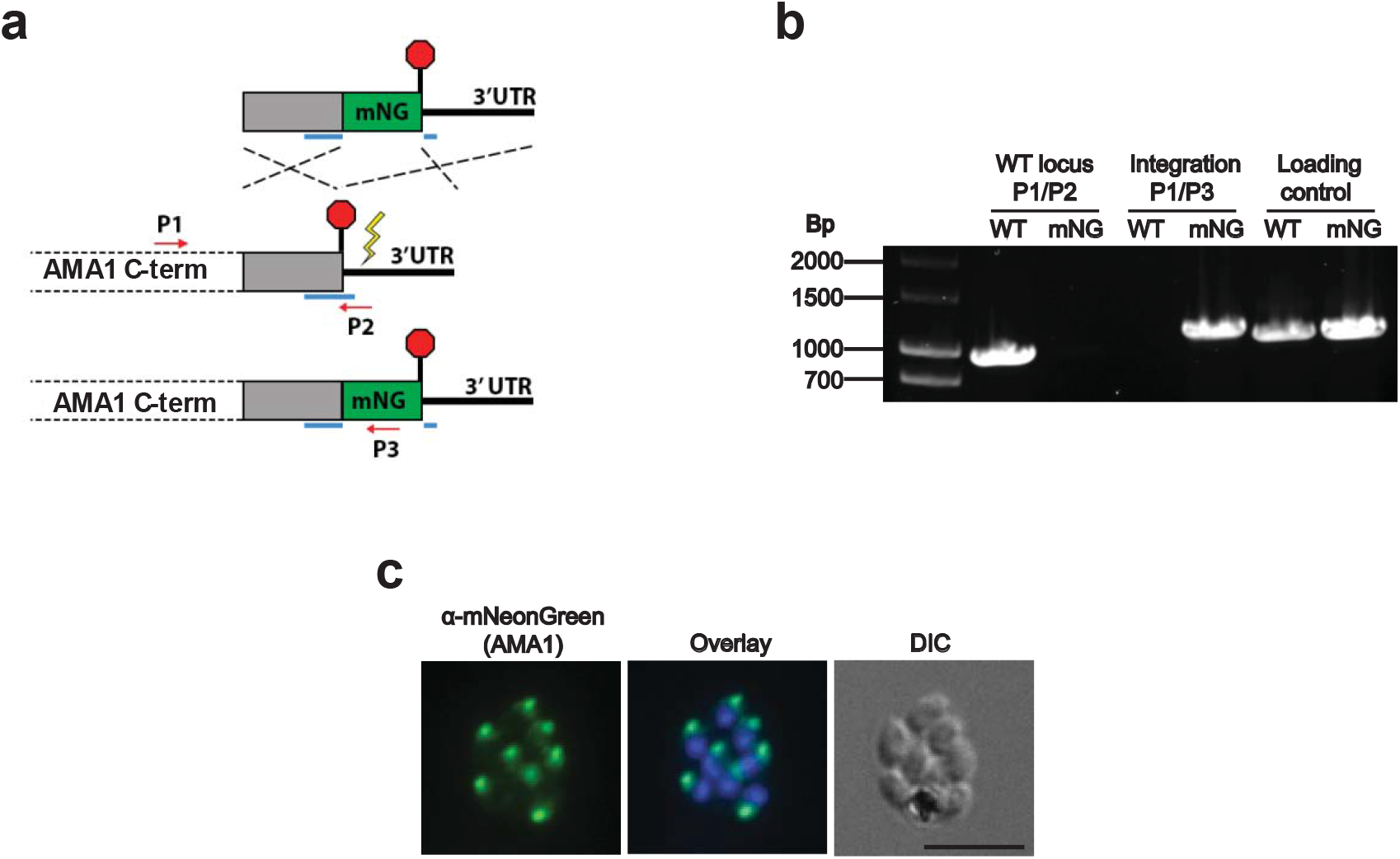
Generation of *Plasmodium knowlesi* parasites with mNeonGreen-tagged AMA1. **a**, Schematic depicting integration of mNeonGreen (mNG)-tagging construct into the *AMA1* locus. Donor DNA was synthesized by overlapping PCR, and consisted of the mNG sequence flanked by 500 bp homologous regions to the C-terminus and 3’UTR of the *AMA1* locus. A sgRNA (position underlined in blue) targeted a CRISPR-Cas9-induced double stranded break (yellow lightning) immediately after the stop codon. Upon repair, the guide sequence was split in two by the insertion of the tag, ablating further Cas9 activity. Positions of diagnostic primers were indicated by red arrows. **b**, Diagnostic PCR showing the absence of wild type (WT) parasites (primers P1/P2; expected band size 988 bp), and the presence of transgenic parasites (mNG; primers P1/P3; expected band size 1118 bp) in the transfected line, along with control PCR reaction detecting unrelated locus. Primers for PCR were listed in Table S1. **c**, Indirect immunofluorescence assay detecting the AMA1-mNeonGreen fusion protein. Antibody specific for the mNeonGreen tag detects the protein expressed in late stage schizonts, which are localised to the apical end of merozoites. Scale bar indicates 5 μm.

**Fig. S3.**
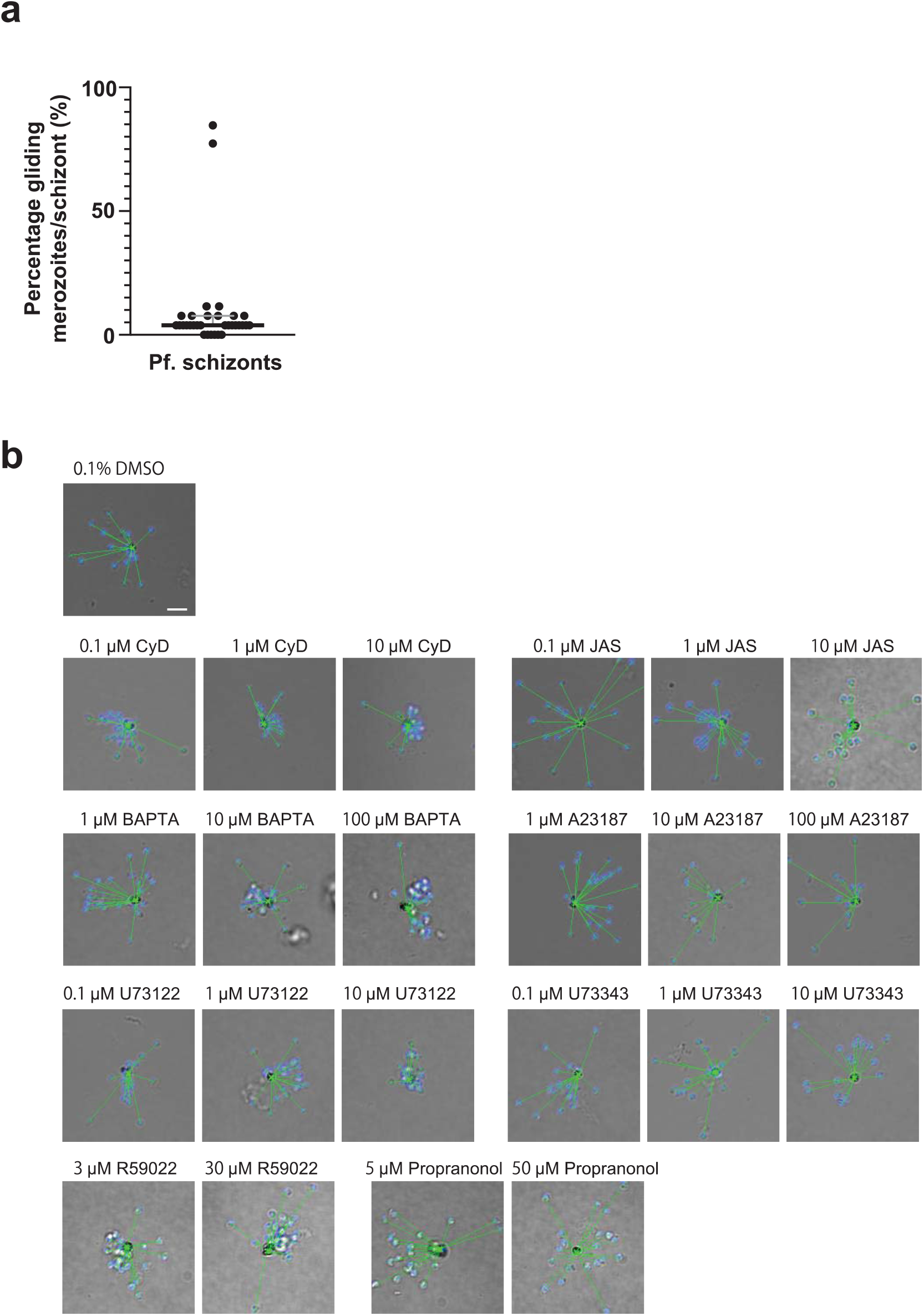
*Plasmodium falciparum* merozoite gliding assay with a panel of chemical compounds. **a**, The percentage of *P. falciparum* merozoites exhibiting motility (n = 32 schizonts). The median (3.85%) and interquartile range are shown. **b**, Compound-treated merozoites were allowed to egress and fixed. The distance between merozoite DNA stained with Hoechst33342 (blue) and hemozoin were measured (green line). Cytochalasin D (CyD), jasplakinolide (JAS), BAPTA-AM (BAPTA), A23187, U73122, U73343, R59022, and propranolol were used in this assay. Scale bar represents 5 μm.

**Fig. S4.**
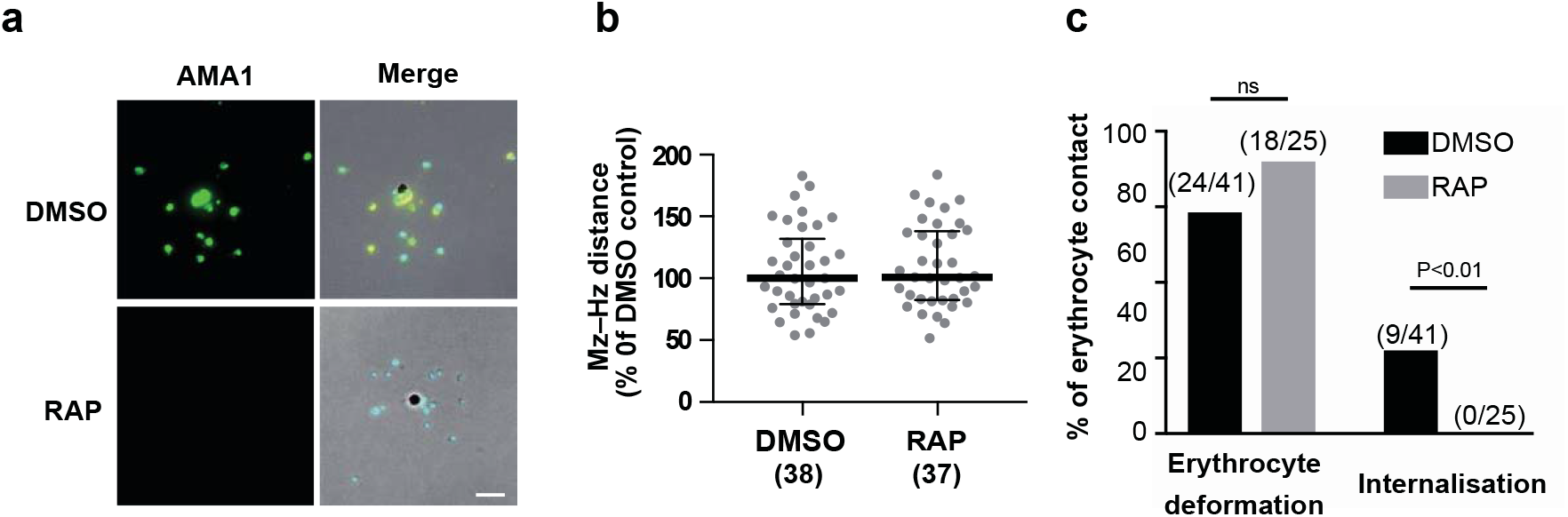
Effect of AMA1-deletion for *Plasmodium falciparum* merozoite gliding motility, erythrocyte deformation, and merozoite internalisation. **a**, PfAMA1:loxP parasite line was treated with DMSO or rapamycin (RAP) and merozoites egressed from infected erythrocytes were stained with anti-AMA1 antibody (green). Right panels, merged images of green AMA1 signals, blue nucleus signals, and differential interference contrast images. **b**, The merozoite (Mz)-hemozoin (Hz) distances (median and interquartile range) were obtained from two biological replicates. No statistically significant difference was detected between DMSO- and RAP-treated parasites by two-tailed Fisher’s exact test. **c**, The number of erythrocyte deformation events was not different between DMSO- and RAP-treated parasites. However, merozoite internalisation events seen for DMSO-treated parasites were not detected in RAP-treated AMA1-deleted parasites (p < 0.01 by two-tailed Fisher’s exact test). ns, not significant. Scale bar represents 5 μm.

### Supplementary Table

**Table S1:**
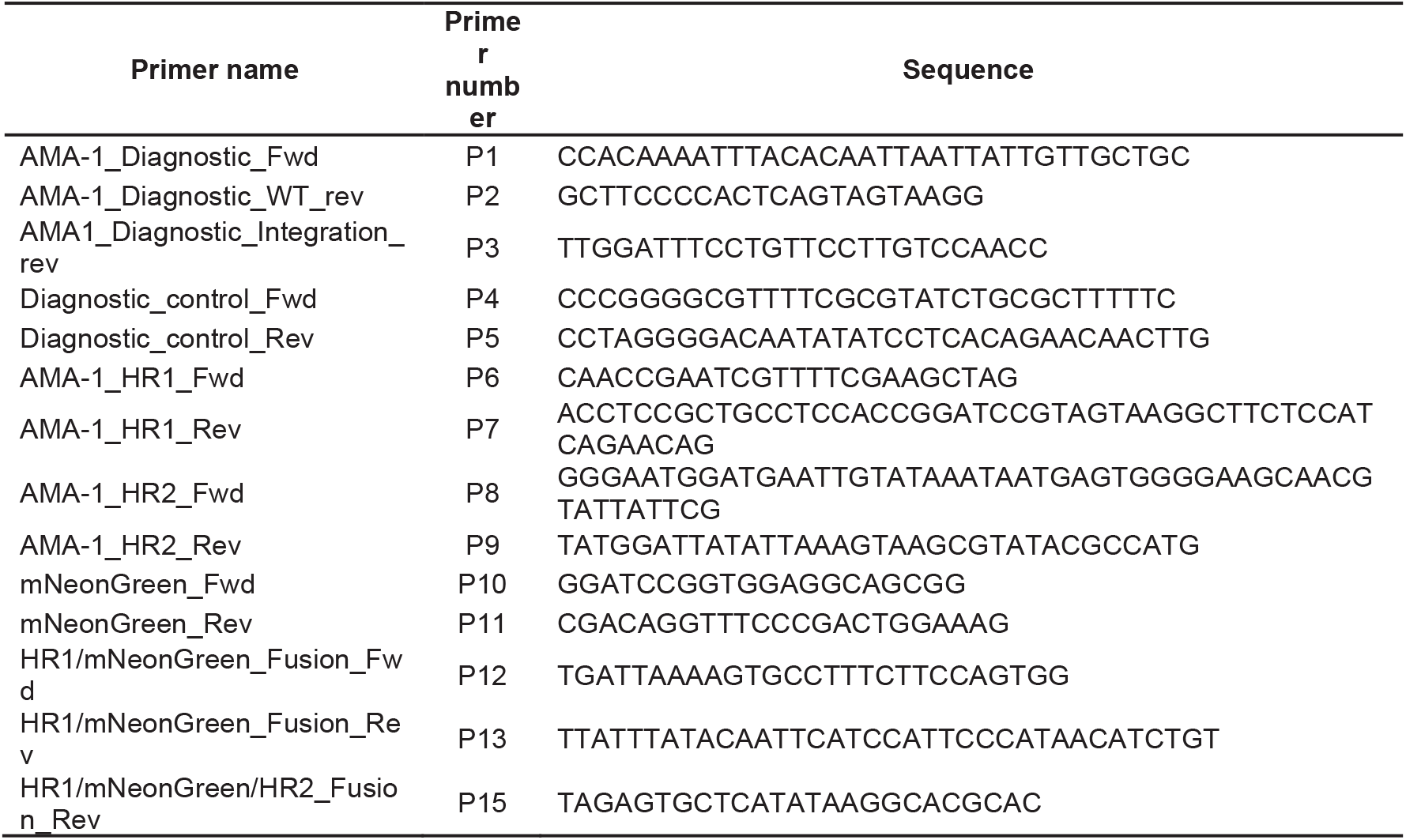
Primers for PCR to generate *Plasmodium knowlesi* parasite with mNeonGreen-tagged AMA1.

### Supplementary Movies

**Movie S1 (separate file).** *Plasmodium falciparum* merozoite gliding motility and erythrocyte invasion on an ibiTreat coverslip (5 frames/second).

**Movie S2 (separate file).** Gliding motility of *Plasmodium falciparum* merozoites with 0.1% DMSO on an ibiTreat coverslip (10 frames/second).

**Movie S3 (separate file).** *Plasmodium knowlesi* merozoites treated with 0.005% DMSO gliding on a poly-L-lysine-coated coverslip (1 frame/second). Parasites were filmed immediately post egress.

**Movie S4 (separate file).** Egress of *Plasmodium knowlesi* merozoites from wheatgerm agglutinin-stained RBCs treated with 100 nM cytochalasin D on a poly-L-lysine-coated coverslip (1 frame/second).

**Movie S5 (separate file).** A *Plasmodium knowlesi* merozoite, designated by a red cross, demonstrating corkscrew-like rotation whilst travelling across a poly-L-lysine-coated coverslip (10 frames/second).

**Movie S6 (separate file).** A *Plasmodium knowlesi* merozoite with mNeonGreen-tagged AMA1 invading an erythrocyte via its ‘wide’ apical end (1 frame/second).

**Movie S7 (separate file).** *Plasmodium falciparum* merozoite gliding motility on an ibiTreat coverslip (10 frames/second). Filming commenced 4 minutes before egress. A red arrow shows merozoite gliding.

**Movie S8 (separate file).** A *Plasmodium knowlesi* merozoite deforming a human erythrocyte whilst gliding across its surface (10 frames/second).

**Movie S9 (separate file).** Egress of *Plasmodium knowlesi* merozoites on human dermal endothelial cells grown on fibronectin-coated μ-Slides (1 frame/second).

**Movie S10 (separate file).** *Plasmodium knowlesi* merozoites completing several short glides on the surface of erythrocytes on a poly-L-lysine-coated coverslip (1 frame/second). A red arrow appears at the beginning of each glide.

